# Identifying Differential Spatial Expression Patterns across Different Slices, Conditions and Developmental Stages with Interpretable Deep Learning

**DOI:** 10.1101/2024.08.04.606512

**Authors:** Yan Cui, Zhiyuan Yuan

## Abstract

Spatially resolved transcriptomics technologies enable the mapping of multiplexed gene expression profiles within tissue contexts. To explore the gene spatial patterns in complex tissues, computational methods have been developed to identify spatially variable genes within single tissue slices. However, there is a lack of methods designed to identify genes with differential spatial expression patterns (DSEPs) across multiple slices or conditions, which becomes increasingly common in complex experimental designs. The challenges include the complexity of cross-slice gene expression and spatial information modeling, scalability issues in constructing large-scale cell graphs, and mixed factors of inter-slice heterogeneity. We propose DSEP gene identification as a new task and develop River, an interpretable deep learning-based method, to solve this task. River comprises a two-branch prediction model architecture and a post-hoc attribution method to prioritize DSEP genes that explain condition differences. River’s special design for modeling spatial-informed gene expression makes it scalable to large-scale spatial omics datasets. We proposed strategies to decouple the spatial and non-spatial components of River’s outcomes. We validated River’s performance using simulated datasets and applied it to identify DSEP genes/proteins in diverse biological contexts, including embryo development, diabetes-induced alterations in spermatogenesis, and lupus-induced splenic changes. In a human triple-negative breast cancer dataset, River identified generalizable survival-related DSEPs, validated across unseen patient groups. River does not rely on specific data distribution assumptions and is compatible with various spatial omics data types, making it a versatile method for analyzing complex tissue architectures across multiple biological conditions.

## Introduction

The advent of spatially resolved transcriptomics technologies has revolutionized our understanding of tissue architectures by enabling gene expression profiling while preserving spatial context^1,2^. As these technologies become increasingly accessible, the scale of experimental data has expanded from hundreds of cells collected from a single slice or a few slices to millions of cells collected from dozens of slices across conditions or temporal stages^3-7^.

This explosion of data has created a pressing need for computational methods that can effectively analyze complex spatial expression patterns of genes at scale^8^. In spatial transcriptomics, a key aspect of spatial data analysis is the identification of spatially variable genes (SVGs), which exhibit significant spatial dependencies in their expression levels^9,10^. SVGs play critical roles in establishing and maintaining tissue organization, and their dysregulation has been implicated in various pathological conditions^11,12^. Several computational methods have been developed to identify SVGs, including early methods like SpatialDE^13^ and Trendsceek^14^, which utilize statistical tests to assess gene spatial variability. Later and recent methods, such as SPARK^15^, SPARKX^16^, SpatialDM^17^, SOMDE^18^, Sepal^19^, and others^20-23^, have improved the accuracy and scalability of SVG identification by introducing various spatial kernels, identifying spatially co-expressed ligand-receptor pairs, employing self-organizing maps, and using diffusion-based processes. Despite these advancements, existing methods primarily focus on identifying SVGs within a single slice. However, with the development of large-scale spatial omics technologies, comparing spatial expression patterns across multiple slices from various conditions has become critical for understanding tissue organization and function.

To address this challenge, we propose a novel task: identifying genes with differential spatial expression patterns (DSEPs) in multi-slice and multi-condition spatial omics data. DSEP genes exhibit changes in spatial expression patterns across different slices, encompassing both gene expression level changes and spatial pattern changes. Existing methods, such as SVG methods and differential expression gene (DEG) methods^24^, are limited in their ability to identify DSEPs, as they focus on single-slice or gene expression abundance without considering spatial information. To overcome these limitations, we developed River, an interpretable deep learning-based method specifically designed to identify genes exhibiting DSEPs among multiple slices and multiple conditions in spatial omics data. River is based on the assumption that only genes with significant DSEPs across slices can contribute to the prediction of slice/condition labels.

We demonstrate River’s performance on carefully designed simulated datasets. We show that River-identified DSEP signal can be decoupled into non-spatial and spatial components. Using a mouse E15.5 embryo dataset, we demonstrate the non-spatial variations of River-identified DSEP genes and validate its generalizability in E16.5 embryos. We also developed a strategy to only pinpoint spatial variation along eight development stages based on a gene expression binarization strategy. In mouse models, River identified diabetes-induced DSEPs in spermatogenesis and lupus-induced DSEPs in the spleen. In human cancer, River identified DSEPs related to Triple-negative breast cancer subtypes, which was validated generalizable across the unseen patient groups. River is also compatible with other spatial omics data other than spatial transcriptomics, for example Multiplexed ion beam imaging by time-of-flight (MIBI-TOF)^25^ and Co-Detection by Indexing (CODEX)^26^. Additionally, River’s special design for modeling spatial-informed gene expression makes it scalable to large-scale spatial omics datasets, making it well-suited for the rapidly accumulating spatial omics data. Our studies demonstrate River’s potential to uncover novel insights into the molecular mechanisms driving spatial heterogeneity and its alterations in different biological contexts.

## Results

### River overview

River (Fig. 1A) is designed to identify genes exhibiting differential spatial expression patterns (DSEPs) among multiple slices in spatial omics data. Each “slice” can be associated with labels such as conditions, developmental stages, disease states, or treatment groups (Fig. 1B). The main idea of River can be summarized as follows: In a multi-slice dataset, DSEPs contribute to differentiate among different slices/conditions, thereby enabling a prediction model to utilize spatially resolved gene expression to distinguish between these slices/conditions. Only genes that show significant changes (spatial and/or non-spatial) between different slices/conditions provide useful information for the prediction model.

**Fig. 1.**
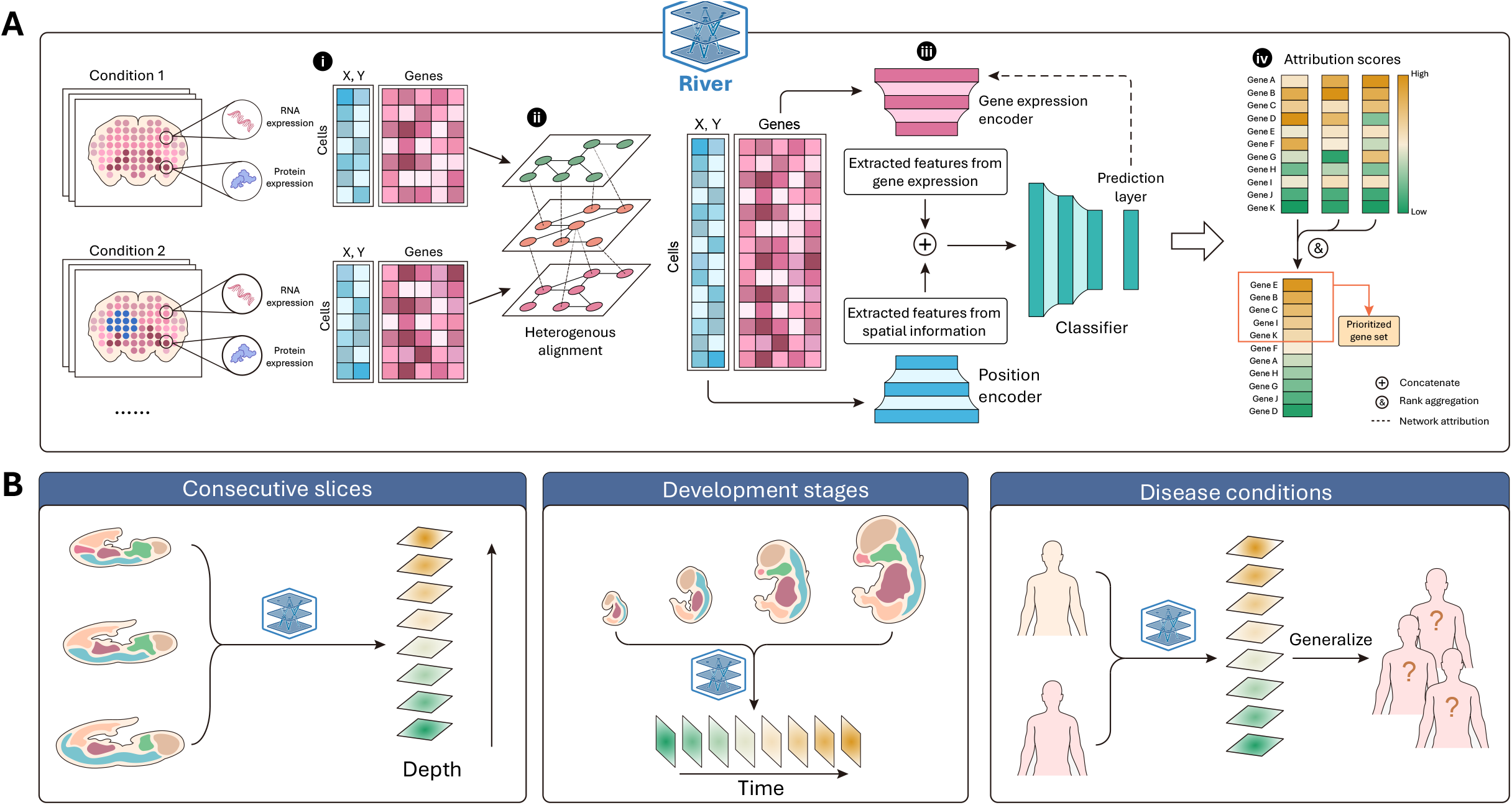
Workflow of River. **A** Workflow of the River process for identifying the differential spatial expression pattern (DSEP) genes in multi-slice and multi-condition spatial omics data, where each slice is annotated with a slice-level label, e.g., development stage or disease. River is composed of two main modules: prediction model and post-hoc attribution. River first fits the prediction model by inputting spatial omics and then utilizes the post-hoc attribution to quantify each gene’s contribution to the prediction. The input spatial omics data for River contains each cell’s expression vector and spatial location, and River utilizes each individual cell’s gene expression incorporated with spatial coordinates as prediction model input. For multi-slice data within different coordinate systems, River applies heterogeneous alignment to ensure a consistent coordinate system for each input cell. River assigns the cell-level input label based on the slice annotation to which it belongs, which is used as supervised information for prediction model training. (i) River utilizes each individual cell as model input instead of using the entire slice as in previous work. River adapts the spatial information by incorporating gene expression with spatial location coordinates in the original slices. (ii) Slices from different spatial coordinates are aligned into the same coordinate space to ensure comparable input spatial coordinates. (iii) The prediction model in River is composed of three parts: position encoder, gene expression encoder, and classifier. The position encoder and gene expression encoder individually encode the input gene expression and spatial coordinates for each cell, obtaining spatial and gene expression latent embeddings, which are concatenated as classifier input spatial-aware gene expression latent. (iv) River utilizes three attribution methods to score the contribution of each gene in the prediction of the target cell-level label based on the fitted prediction model. The outcome of this tentative attribution process is three independent gene score vectors. River utilizes rank aggregation to combine the multiple attribution results to form a final gene rank list. **B**, River can be utilized in diverse multi-slice scenarios, including consecutive slices, development stages, and disease conditions.

River (Fig. 1A) is based on interpretable deep learning, consisting of a prediction model followed by deep learning attribution methods to identify the genes contributing to the prediction. These contributions are quantified as scores to prioritize DSEP genes. The process can be broken down into the following steps: (1) designing the prediction model to fully utilize the spatial-aware gene expression features in a multi-slice and multi-condition dataset, and (2) quantifying contributions of each gene to the prediction model.

To aggregate spatial position and gene expression data (Fig. 1A-i) and obtain spatial-aware gene expression latent representations for each input cell, River utilizes a joint two-branch architecture. This architecture includes a position encoder (to extract features from spatial information) and a gene expression encoder (to extract features from gene expression), which independently extract features and then fuse them in the latent space (Fig. 1A-iii). Before input into position encoder, cells from different slices are spatially aligned using heterogeneous alignment methods to harmonize spatial information (Fig. 1A-ii). Above approach ensures that gene expression features in the latent space are spatial information-aware, which are then sent to subsequent modules to predict slice-level labels. The position encoder is motivated by its efficiency and scalability compared to previous graph neural network based spatial embedding methods, while maintaining information integrity, as demonstrated in previous spatial omics studies^27^. After the training phase, River employs multiple deep learning attribution strategies to obtain cell-level gene contribution scores, which are then aggregated to derive the final global scores (Fig. 1A-iv). Compared with direct and global-level selection methods like Lasso^28^, the instance-level scores reflect the high cell-wise heterogeneity in spatial omics slices. A rank aggregation method synthesizes the contribution rankings provided by multiple interpretation attribution techniques. This aggregation process is critical, as it combines insights from multiple analytical perspectives and obtains a robust and reliable measure of each gene’s contribution on the prediction of slice labels. More details can be found in Methods.

Both the training of the prediction network and the attribution module can be conducted in mini-batch distributed computation on GPUs. Moreover, the use of non-graph spatial information embedding techniques ensures the scalability and efficiency of River on large-scale multi-slice data, which is critically important in modern large-scale biological studies.

### Benchmarking analysis

To evaluate the performance of River, we generated simulated datasets (Fig. 2A, see Methods). The control slice (slice 0) contained four different spatial domains. Condition slices (Slices 1 – 6) were generated based on slice 0, each with a carefully designed and distinct perturbation to control gene expression variability (Fig. 2A). This setup provided a ground truth where perturbed genes were labeled as positive (DSEP genes) and the remaining genes as negative (background or non-DSEP genes), which allowed us to evaluate the performance of various methods. We compared slice 0 with each of slices 1 – 6 (resulting in datasets 1 to 6) using different methods (Fig. 2B).

**Fig. 2.**
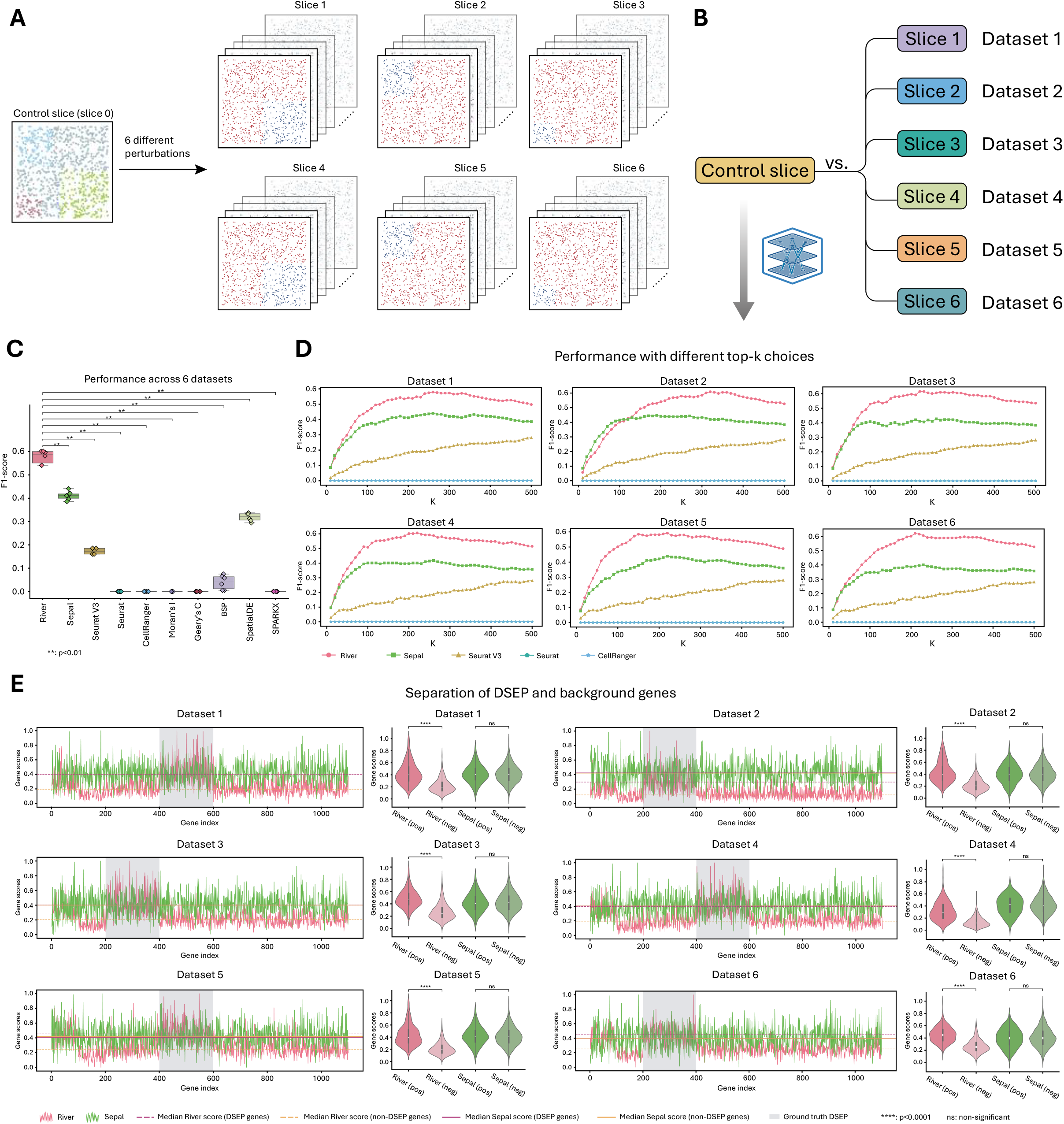
Simulation benchmarking. **A**, 6 different perturbations were applied on the control slice (slice 0) in silico, obtaining 6 new slices (details in Methods). **B**, The control slice (slice 0) was compared with each of slice 1 to slice 6 using River. **C**, Benchmarking outcome for each method on six datasets. The performance of different methods is evaluated by F1-scores. River achieves the highest F1 score across six experiments with statistical significance (p value < 0.05, rank-sum test). **D**, Benchmarking results summary for top k parameter dependency methods among different k values in F1 scores. X-axis: different k choices. Y-axis: F1-score. **E**, Comparison of score distribution between River and Sepal. River’s attribution method is IG for this figure, other two methods are also compared with Sepal in Supplementary Fig. 2. For each dataset, the left line chart indicates the score value for each gene, where positive genes (Ground truth DSEP genes) are expected to obtain larger scores compared with remaining negative genes. The right violin plot indicates the score distribution for the two methods between the DSEP and non-DESP genes. P-values are obtained using rank-sum test.

Since there are currently no methods for identifying differential spatial expression pattern (DSEP) genes across slices, we adapted existing methods to be compatible with the multi-slice context. Specifically, our competing methods included adapted highly variable gene (HVG) selection methods (SeuratV3^29^, Seurat^30^, CellRanger) and adapted spatially variable gene (SVG) selection methods (SPARKX^16^, SpatialDE^13^, Sepal^19^, Moran’s I^31^, Geary’s C^31^, and BSP^32^). We explained how these methods can be adapted to multi-slice analysis (see Methods).

The performance of River and nine competing methods was summarized across six datasets (Fig. 2C). River significantly outperformed all other methods in terms of F1-score (p-value < 0.05). River ranked first with a median F1-score of around 0.59, while the second and third best methods, Sepal and SpatialDE, had median scores of approximately 0.41 and 0.32, respectively (Fig. 2C). The other methods had F1-scores close to zero, indicating their inadequacy for this challenging task (Fig. 2C). Additionally, since River, Sepal, SeuratV3, Seurat, and CellRanger can output gene-wise scores, we compared the F1-scores using different top-k choices for these methods (Fig. 2D). Regardless of the selection of k, River outperformed the other methods in almost all cases (Fig. 2D).

River’s attribution module (see Methods) can output meaningful scores for each gene, prioritizing those with differential spatial expression patterns. To further validate River’s attribution scoring capability, we analyzed whether River’s attribution score could differentiate true DSEP genes from background genes (Fig. 2E). We compared one of the attribution methods (see Methods), Integrated Gradient (IG) score of River with Sepal (second best method that can output gene-wise scores as shown in Fig. 2C-D). For each dataset, we plotted the gene-wise scores provided by River and Sepal (Fig. 2E). Across the six datasets, River (represented by the red curve) consistently assigned higher scores to DSEP genes compared to background genes (red dashed lines always higher than orange dashed lines across 6 datasets), with the scores exhibiting significant differences between true DSEP genes (denoted as River (pos)) and background genes (denoted as River (neg)) (p-value < 0.05) as shown in the violin plots (Fig. 2E). In contrast, Sepal (represented by the green curve) failed to differentiate DSEP genes from background genes (red solid line always overlap with orange solid line), as Sepal-computed scores did not show significant differences between the two groups, i.e., Sepal (pos) and Sepal (neg), as shown in the violin plots (Fig. 2E). Additional comparisons of other attribution methods can be found in Supplementary Fig. 2.

### River detects non-biological spatial expression patterns across slices

When comparing slices, differential spatial expression patterns (DSEPs) identified by River can arise from both spatial and non-spatial variations. Non-spatial variations may originate from gene expression level differences between slices, either due to biological or non-biological factors (e.g., batch effects). We used a mouse embryo dataset^33^ to demonstrate River’s capability to identify genes with non-spatial variations among slices. Specifically, this dataset contains four replicate slices of E15.5 mouse embryos and another four replicate slices of E16.5 mouse embryos. (Fig. 3A)

**Fig. 3.**
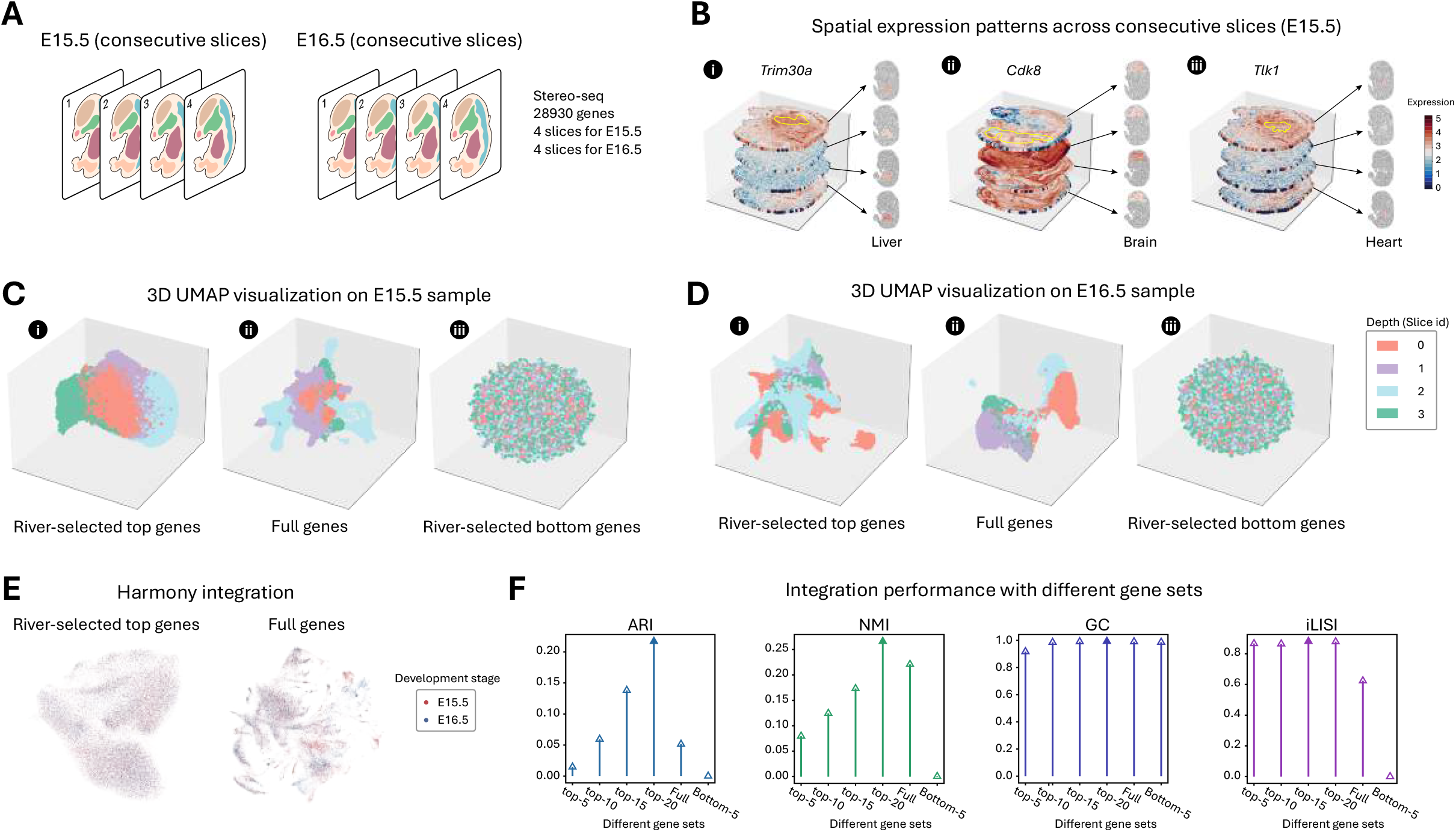
Analysis of Stereo-seq mouse embryo datasets E15.5 and E16.5. **A**, Dataset: Stereo-seq mouse embryo dataset in E15.5 and E16.5 development time, each with four continuous depth slices. River utilizes the depth as the slice level label and fits the model on the E15.5 dataset. **B**, 3D visualization of the spatial gene expression pattern of River-selected top-3 genes at the E15.5 timepoint: *Trim30a, Cdk8*, and *Tlk1*. **C**, 3D UMAP visualization of River-selected top-20 genes, full gene panels, and River-selected bottom-20 genes on E15.5 timepoint data for each cell. Points in the figure are colored by each cell’s slices id. **D**, 3D UMAP visualization of River-selected top-20 genes (fitted in E15.5), full gene panels, and River-selected bottom-20 genes on E15.5 timepoint data for each cell. Points in the figure are colored by each cell’s slice id. **E**, 2D UMAP visualization of Harmony integrated datasets (development stage as batch key) used in Fig. 3A, using River-selected top genes (left) and full genes (right). **F**, Batch integration metrics comparison among different River-selected top-k (k=[5,10,15,20]). River-selected genes with bottom genes and full genes. The metrics include ARI, NMI, Graph Connectivity (GC) and iLISI.

We applied River to detects genes that can differentiate the four slices of the E15.5 dataset. Since these slices were consecutive from the same embryo, any differential genes among them were likely attributed to non-biological factors (e.g., caused by different experimental batches or z-axis differences) (Supplementary Fig. 3A). Visualizations of the top-3 River-identified genes (*Trim30a, CDK8*, and *Tlk1*) in a stacked 3D space confirmed their distinct expression patterns across the slices (Fig. 3B). Specific regions, including liver (Fig. 3B-i), brain (Fig. 3B-ii), and heart (Fig. 3B-iii), highlighted the different expression levels evidently.

To validate that River-prioritized genes can be attributed to non-biological factors, we performed Uniform Manifold Approximation and Projection (UMAP)^34,35^ on all cells from the four slices using different gene sets (full gene set, River-selected top-20 genes, and River-selected bottom-20 genes) as input features to assess the information contained in River-selected genes and their negative controls. When using the top-20 genes ranked by River, cells from each slice clustered together and were clearly separated in the 3D UMAP space (Fig. 3C-i), indicating that these genes effectively captured the most prominent differences of the consecutive slices. Using all genes as features resulted in a less distinct separation of the slices (Fig. 3C-ii). On the negative control, using the River-selected bottom-20 genes as features completely failed to distinguish the slices (Fig. 3C-iii), confirming that these genes had the lowest inter-slice differences.

We hypothesize that these top-ranked genes are also most affected by non-biological factors like batch effects in other samples. To test the generalizability of the River-selected gene sets, we applied the top-20 genes identified in the E15.5 slices to four slices from the E16.5 embryo. Strikingly, the 3D UMAP using these genes effectively separated the E16.5 slices (Fig. 3D-i), demonstrating the robustness of River-identified genes on different animal. The UMAP using all genes in the E16.5 slices can also separate different slices (Fig. 3D-ii). And the bottom-20 genes (negative control) from E15.5 failed to distinguish the E16.5 slices (Fig. 3D-iii).

We hypothesize that these genes can be used to improve data integration. By applying Harmony^36^ to the top River genes, we observed a superior mixing of cells from E15.5 and E16.5 in the UMAP space compared to using all genes (Fig. 3E). We used the well-known integration benchmarking pipeline, scib^37^, to evaluate the integration performance using different gene sets (see Methods). The results confirmed that the top genes identified by River can substantially improve the data integration (Fig. 3F).

In the above analyses, we did not try to avoid batch effects but used these signals as a sanity check to demonstrate that River can identify different gene signals across slices. We also showed that such batch effect genes exhibit similar behaviors in other samples and demonstrated the improvement in downstream data integration analyses.

### River uncovers DSEP genes across developmental stages

To demonstrate the diverse and generalized utility of River in identifying DSEP genes across different conditions in multi-slice datasets, we focus on another spatial omics application of interest to the research community: temporal changes^38^. Existing studies often focus on gene spatial patterns within the same slice, overlooking changes in spatial gene expression patterns over time. Here, we applied River to the Stereo-seq dataset of mouse embryos spanning eight development stages^33^ (same sectioning position in respective animals) (Fig. 4A). In this case, River-identified differential genes may be attributed to both spatial and non-spatial variations caused by development.

**Fig. 4.**
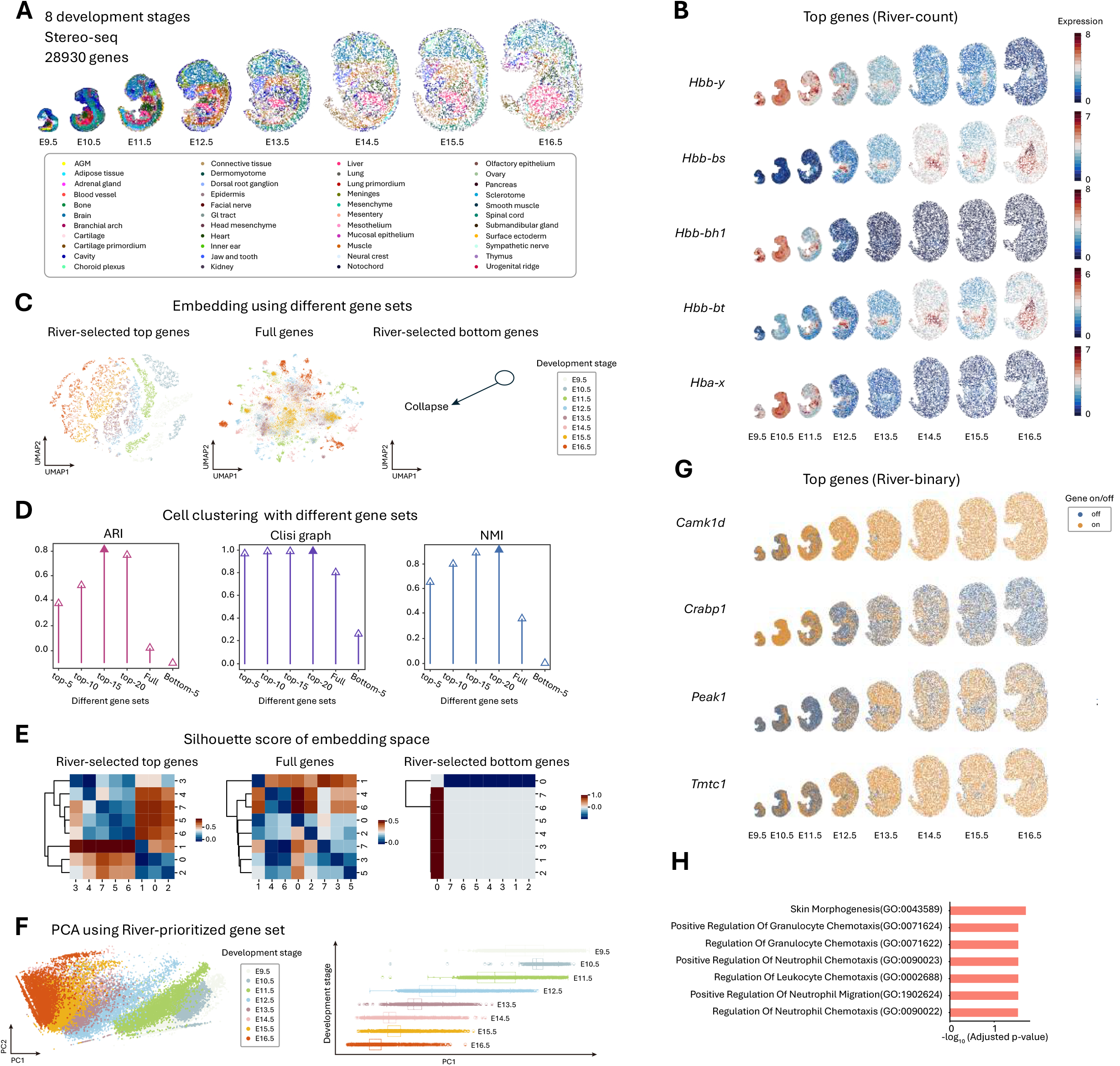
Analysis of Stereo-seq mouse embryo datasets with 8 development stages. **A**, Dataset: Stereo-seq mouse embryo dataset across eight development stages ([E9.5, E10.5, E11.5, E12.5, E13.5, E14.5, E15.5, E16.5]). **B**, Visualization of River-identified top-5 genes using count expression values. **C**, t-SNE visualization of different gene set inputs (River-selected top-5 genes, full genes, and River-selected bottom-5 genes). Notably, the bottom-5 genes’ t-SNE is collapsed in the t-SNE embedding space due to the lack of information, providing a negative control for River-selected genes. **D**, Unsupervised clustering results comparison between different input gene sets (top-5 genes, full genes, and bottom-5 genes) using NMI, ARI, and cLISI metrics. **E**, Pairwise silhouette score across eight development stages for different input gene sets. The pairwise silhouette score reflects the distance between two clusters in the t-SNE space in Fig. 4C. **F**, Principal Component Analysis for River-selected top-5 genes colored by development stages. The right panel shows the principal components’ distribution change tendency along with developmental changes. **G**, Visualization of River-identified top genes using binary expression values (unique to the gene set of River-count). **H**, Significant enriched (FDR adjusted p value < 0.05) gene sets uniquely identified by River-binary in GO biological process reference.

Visualization of the top-5 genes identified by River confirmed their spatiotemporal variation along the developmental axis (Fig. 4B). To assess the effectiveness of the top genes in distinguishing different developmental stages, we performed t-distributed Stochastic Neighbor Embedding (t-SNE)^39^ using the top-5 genes identified by River. We found that the embedding space effectively separated cells from different stages (Fig. 4C left), with visually better separation compared to using all genes (Fig. 4C middle). In contrast, using the bottom five genes completely failed to distinguish the stages (Fig. 4C right), as the t-SNE visualization collapsed due to the gene expression value similarity. This provides a strong negative control example for the River-selected gene set. These findings were further supported by three quantitative metrics (NMI, ARI, and Cell-type LISI (cLISI) score, see Methods) on different gene set-constructed embedding spaces, confirming that the top-5 genes selected by River contain the non-spatial variations (gene expression level, Supplementary Fig. 3B) to discriminate across slices (Fig. 4D).

Furthermore, we used the pairwise silhouette score to measure the distance similarity for each time point in the input feature space obtained using different input gene sets. Fig. 4E shows that in the River-selected top-5 input gene set, the closer time points share more similar pairwise silhouette score patterns and show better clustering compared to the entire input gene set, indicating that the non-spatial variation captured by River contains biological signals related to development. However, it has been reported that t-SNE retains local data structure better than global data structure^40-43^, meaning that the cell group distance (global structure) in the t-SNE embedding recorded in the silhouette score may be blurred. To more strictly test the biological relevance of River-prioritized genes, we performed Principal component analysis (PCA), which retains global data structures, using the top-5 River genes. PCA using only the top-5 genes identified by River showed clear separation of cells from different stages, indicating that these top genes successfully captured non-spatial gene expression variations (Fig. 4F left). Interestingly, we observed that the slices arranged in a consistent order along the developmental timeline in the PCA space (Fig. 4F left), and PC1 alone significantly separated the different stages in the correct order (Fig. 4F right), indicating that these non-spatial gene expression variations are not solely due to non-biological factors such as batch effects and indeed contain biological signals related to development.

To demonstrate River’s capability to capture spatial pattern differences (Supplementary Fig. 3C), we decoupled the spatial and non-spatial variations using a gene expression binarization approach (see Methods). This process transforms all gene expression in the input slices into 0/1 values (0 for off and 1 for on), and we used this binarized expression as input for River. The benefits of this procedure are that (1) binarized gene expression is reported to be robust to batch effects in both single cell^44,45^ and spatial transcriptomics^46,47^, and (2) binarized gene expression removes gene expression level variations and only retains spatial patterns. River with binary input (River-binary) identified the top-10 ranked pure spatial pattern shift genes, and six of these were the same as those identified using the count expression value-informed River (River-count) outcome (*Hbb* family genes, Fig. 4B). This demonstrated River’s ability to capture spatial variations. We present the remaining four uniquely selected genes from River-binary results, all of which show significant spatial pattern shifts across the developmental stages (Fig. 4G). We further conducted gene set enrichment analysis for the gene set selected by River-binary (pure spatial pattern shift) uniquely to River-count. The results showed that these unique genes are highly enriched in two main biological processes occurring during embryo development: chemotaxis and skin morphogenesis, indicating that River can capture development-related spatial pattern changes (Fig. 4H).

### River identified diabetes-induced DSEP genes in spermatogenesis

Apart from applications in multi-slice developmental studies, River can also be used to analyze spatial properties in tissue samples from both normal and diseased states. This capability is crucial for uncovering complex cellular interactions and gene expression patterns associated with disease mechanisms. To illustrate this, we applied River to study the impact of diabetes on spermatogenesis in mice. The input dataset comprised testis sections from three wild-type (WT) and three leptin-deficient diabetic mice^48^ (Fig. 5A). River utilized the WT and diabetic annotations for each slice as labels during model fitting. The top-ranked genes selected by River are displayed in Fig. 5B. Notably, *Prm1* and *Prm2*, previously associated with ES/spermatozoon loss in diabetic testes^49^, were identified by River.

**Fig. 5.**
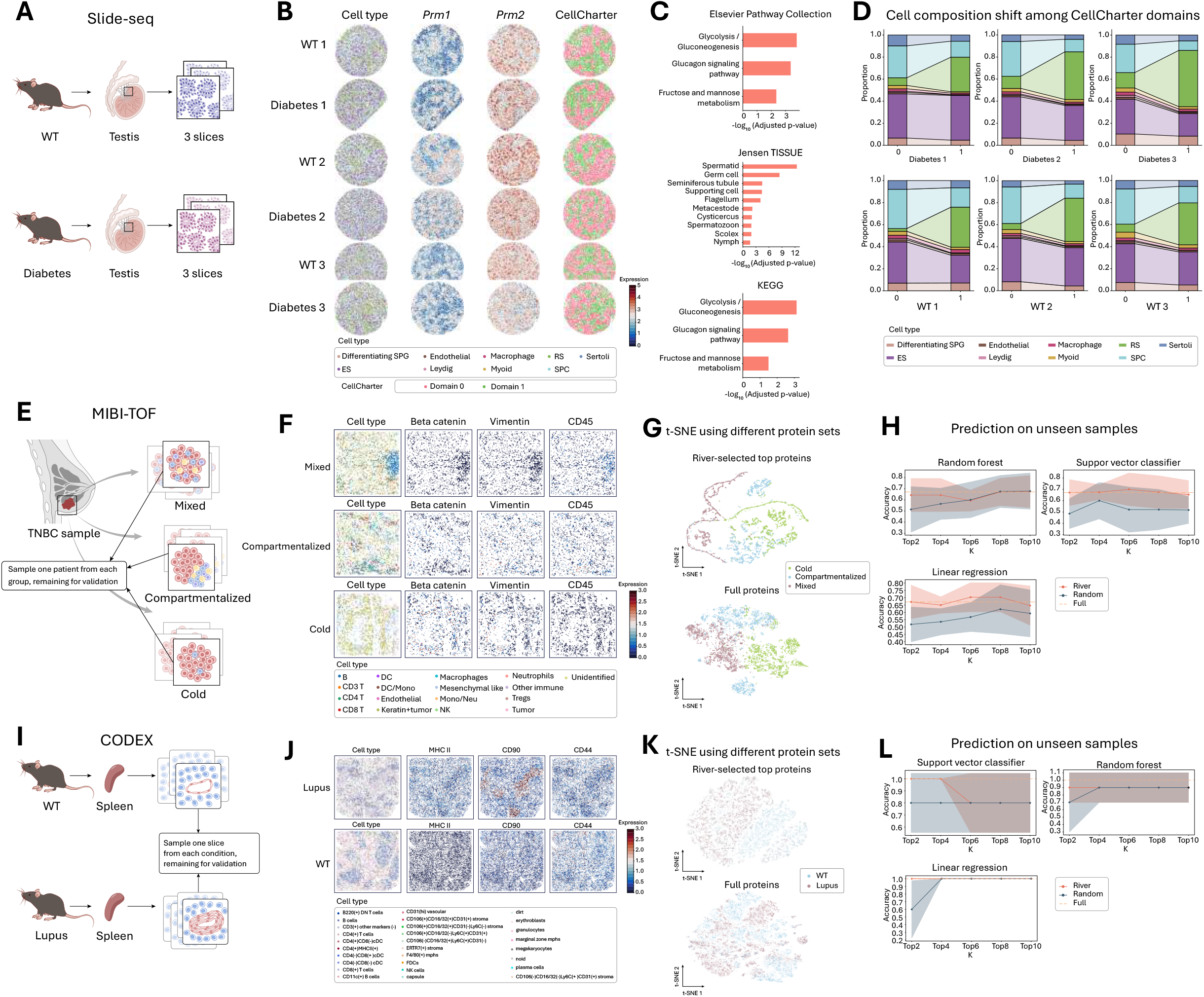
Applications in 3 disease cases (Slide-seq, MIBI-TOF, and CODEX datasets). **A**, Slide-seq Dataset: 6 mouse testis Slide-seq slices (3 Diabetes and 3 WT). River utilized the Diabetes/WT condition label to select the diabetes-induced pathological changes related genes. **B**, The visualization of cell type, spatial expression gene pattern of River-selected top-ranked genes (*Prm1, Prm2*), and CellCharter co-clustering results based on River-selected top-200 genes for input 6 slices. **C**, Significantly enriched gene sets (FDR adjusted p-value < 0.05) in gene set enrichment results for River-selected top-50 genes on three reference gene sets: KEGG, Jensen TISSUE, and Elsevier Pathway Collection. **D**, Cell composition shift between CellCharter identified domain 0 and domain 1 in each of the six individual slices. Round spermatids (RSs) and spermatocytes (SPCs) composition. **E**, MIBI-TOF Dataset: MIBI data from 41 patients with 19 Mixed (high immune infiltration), 15 Compartmentalized (distinct tumor and immune cell regions), and 6 Cold (low immune cell presence). We randomly chose one sample per category to fit River. Remaining hold-out slices were utilized for the validation of River-selected biomarker panel generalization. **F**, Visualization of River-selected top-3 panel, showing significant spatial expression pattern shifts across three TNBC subtypes. **G**, t-SNE visualization of cells using the River-selected top-5 panel and full original panel as input features. Cells are colored according to patient label. **H**, The validation results of the River-selected panel generalization on hold-out slices. We conducted 5-fold validation on the hold-out slices for three baseline classifiers (Support Vector Classifier, Random Forest, and Logistic Regression) with different top-k parameters in both River, randomly chosen baseline, and full panel. The line chart shows the 5-fold accuracy on the validation set with mean and confidence interval for different k ([2, 4, 6, 8, 10]). **I**, CODEX Dataset: CODEX data from 9 mice with 3 WT and 6 lupus spleens. We randomly chose one sample per category to fit River. Remaining hold-out slices were utilized for the validation of the River-selected biomarker panel generalization. **J**, Visualization of the River-selected top-3 panel and cell type, showing significant spatial expression pattern shifts across WT and lupus, and high relevance to specific cell types (e.g., CD90 with T-cells). **K**, t-SNE visualization of cells using the River-selected top-5 panel and full original panel as input features. Cells are colored according to patient label. **L**, The validation results of the River-selected panel generalization on hold-out slices. We conducted 5-fold validation on the hold-out slices for three baseline classifiers (Support Vector Classifier, Random Forest, and Logistic Regression) with different top-k parameters in both River, randomly chosen baseline, and full panel. The line chart shows the 5-fold accuracy on the validation set with mean and confidence interval for different k ([2, 4, 6, 8, 10]).

To ensure robust results, we performed gene set enrichment analysis on the top-50 genes identified by River using three reference gene sets: KEGG, Jensen TISSUE, and Elsevier Pathway Collection (https://maayanlab.cloud/Enrichr/) (Fig. 5C). The enrichment results indicated that, from both pathway and tissue composition perspectives, the River-selected genes were significantly enriched in diabetes-induced pathological changes in spermatogenesis. Specifically, the Elsevier pathway analysis showed enrichment in the male infertility pathway, while KEGG analysis revealed enrichment in the Glycolysis/Gluconeogenesis pathway, aligning with previously reported disturbances in the male reproductive system associated with diabetes^50^. In terms of tissue composition, the Jensen enrichment results demonstrated significant enrichment of River-selected genes in spermatogenesis-related tissue components in the testis, such as spermatids, germ cells, and seminiferous tubules. This supports that River can capture genes pertinent to spermatogenesis-related cells and tissue composition.

To further validate the relevance of these genes to cell and tissue composition, we utilized the prioritized genes as input features for CellCharter^27^ to conduct multi-slice spatial co-clustering and identify consistent tissue compositions across all slices in the dataset (see Methods, Multi-slice Spatial Co-Clustering). The clustering results showed that *Prm1* and *Prm2* shared a similar spatial arrangement within the identified domain arrangement, indicating consistent spatial expression patterns among the top-ranked genes (Fig. 5B). This suggests that River-selected genes can be used to identify continuous tissue compositions across slices. Additionally, we compared cell type compositions between domains 0 (red) and 1 (green) in each slice identified by CellCharter (Fig. 5D). The early round spermatids (RSs) and spermatocytes (SPCs) exhibited the most significant shift among the input slices (Fig. 5D). This shift reflects one of the spermatogenesis processes where spermatocytes divide to form round spermatids^51^, suggesting that River-identified diabetes-induced pathological change-related genes reveal tissue compositions related to spermatogenesis.

### Applications on spatial proteomics datasets

We applied River to two spatial proteomics datasets to test its generalization potential on platforms other than spatial transcriptomics. First, we used a triple-negative breast cancer (TNBC) spatial proteomics dataset^52^ measured by MIBI-TOF. The dataset featured three patient groups associated with significantly different survivals: Mixed (high immune infiltration), Compartmentalized (distinct tumor and immune cell regions), and Cold (low immune cell presence) (Fig. 5E). For each patient group, we selected one patient to train River and used the River score to prioritize the protein set. We visualized the top-ranked proteins— Vimentin, Beta-catenin, and CD45—on each patient in Fig. 5F. Each of these proteins exhibited distinctive spatial patterns across patients. For instance, CD45, a marker of immune cells, showed denser expression patterns in both Mixed and Compartmentalized conditions. Meanwhile, Beta-catenin and Vimentin, markers of TNBC tumor cells, displayed a more scattered distribution in Mixed conditions and a dense, centralized distribution in Compartmentalized conditions, reflecting the characteristics of immune infiltration and separation^52^. Additionally, we performed t-SNE visualizations for the top-5 proteins identified by River, showing similar separation than using the full protein set (Fig. 5G).

Despite the original study’s efforts to handle batch effects in MIBI-TOF data generation (Supplementary Fig. 4A), batch effects might still persist. To validate the biological significance of the signals captured by River, we tested whether River-identified DSEPs contain biological variations applicable to unseen patients. We employed the genes ranked in the different top-k (k=[2, 4, 6, 8, 10]) identified on one patient and tested on unseen patients. This selected protein set was used to compute the mean expression value as a patient-level feature. We then analyzed this feature using three classifiers (Support Vector Classifier (SVC), Linear Regression (LR), and Random Forest (RF)) with 5-fold validation. The results, shown in Fig. 5H, indicate that the selected genes maintained comparable predictive power to the original full gene set and remained robust to the selection of k parameter, with consistent improvements compared to the random selection.

Next, we tested River on a second dataset. This dataset contains spatial proteomics (CODEX) data measured on lupus spleen mouse model with two condition labels: WT and lupus^26^ (Fig. 5I). We followed the same setup as the previous TNBC dataset, choosing one slice from each condition to train River and select top proteins. The visualizations of the River-selected top-ranked proteins (CD44, MHC class II, CD90) abundance on the chosen slice are shown in Fig. 5J. Previous studies have identified MHC class II as associated with lupus susceptibility^53^. Specifically, in mice, the MHC class II locus directly contributes to lupus disease susceptibility, similar to observations in humans. Additionally, CD44, a surface marker of T-cell activation and memory, was overexpressed in T cells of lupus spleen^54^. CD90, a marker of T cells, also showed significant changes in the T cell niche between lupus and WT conditions^55^. These proteins reflect the properties of lupus at both molecular and cell niche interaction levels.

We repeated the visualization procedure from the previous TNBC experiment, performing t-SNE visualizations for the full protein panel and the River-prioritized proteins (Fig. 5K). To further validate the predictive power of the River-selected panel on hold-out unseen slices, we conducted quantitative prediction tasks. Following the same settings as the TNBC experiment, we trained SVC, LR, and RF using the mean value of the chosen panel for different top-k (k=[2, 4, 6, 8, 10]) in the hold-out unseen slices with five-fold validation. The results showed that River-prioritized proteins obtained comparable accuracy with the full protein panel, better than random baseline (Fig. 5L).

### Scalability and reproducibility

River’s non-graph design greatly enhances its scalability for handling large-scale spatial datasets. Contrary to most existing methods that rely on graph structures to model cell-cell spatial relationships and suffer from scalability issues with large graphs (e.g., when a slice contains a vast number of cells), River efficiently manages such challenges. We demonstrated this using a brain spatial transcriptomics dataset measured by MERSCOPE (https://vizgen.com/resources/using-merscope-to-generate-a-cell-atlas-of-the-mouse-brain-that-includes-lowly-expressed-genes/), which contains three replicates (Supplementary Fig. 1A). Each slice comprises more than 70,000 cells, posing significant computational hurdles for many existing methods, as previously highlighted^56-58^. However, River processes each replicate in ∼ 7 minutes (machine information in Methods), showcasing its efficiency. Furthermore, the three replicates allowed us to test River’s reproducibility, and we observed that River consistently assigned gene-wise scores across the replicates (Supplementary Fig. 1B).

## Discussions

Scalable spatial omics technologies have enabled the generation of large-scale multi-slice and multi-condition datasets. One key insight from such datasets is the identification of differences in spatial gene expression patterns across different conditions, which had previously been overlooked. We propose a new concept: Differential Spatial Expression Patterns (DSEPs). DSEPs refer to changes in a gene’s spatial expression pattern across different slices or conditions, encompassing changes in spatial arrangement, gene expression level, or both. This concept is more suitable for characterizing gene spatial properties in large-scale multi-slice studies than previous concepts, such as Differential Gene Expression (DEG) analysis and Spatially Variable Genes (SVGs) analysis.

We developed River, a method that uses interpretable deep learning to identify DSEPs across slices. Our results demonstrate that River can effectively identify DSEPs genes across extensive multi-slice and multi-condition spatial omics datasets, making it the first method to do so at scale. River is not simply another differential gene expression or SVG identification method but is specifically designed to identify DSEPs without being limited by single-slice and cell-independent hypotheses. Furthermore, we have demonstrated River’s biomedical significance using various biological cases such as development and disease, which cannot be done with previous methods.

River’s novel point, which transforms the differential spatial expression pattern identification problem into a solvable computational task with interpretable deep learning, holds potential for future studies, especially those aiming to uncover factors significantly contributing to certain condition labels. This includes identifying cell states responding to certain perturbations and pinpointing microenvironments exclusive to certain diseases^59-61^. Additionally, River’s use of non-graph data structures to model cell-cell spatial relationships offers valuable insights for future spatial omics data modeling.

Several areas for future improvement remain. One major concern in comparing different slices and conditions is the batch effect. In this study, we eliminated this using two approaches: (1) a gene expression binarization method and (2) utilizing batch-effect-free datasets pre-processed by the original studies. Future research could enhance this framework by incorporating contrastive modules to create an end-to-end solution. Another potential enhancement involves using single-cell foundation models^62-66^ to replace River’s gene expression encoder, known for their robustness against batch effects. In cases where different slices and conditions originate from various spatial platforms and resolutions, employing recently proposed rasterization techniques^67,68^ could be beneficial. This would allow the direct comparison of data from different resolutions within the same spatial framework and make the analysis scalable to very large-scale datasets.

## Method

### Overview of River

River can be considered as a combination of two main functional modules. The first module is the prediction model, which utilizes spatial omics (transcriptomics/proteomics and other modalities; for simplicity, we refer to the input feature as gene expression in the following content) features and spatial location (represented by spatial coordinates) for each single cell as input. This module is made up of a Multi-Layer Perceptron (MLP) due to its representative capability. The training target of the model is predicting the condition label for each input cell, defined by the corresponding original slice-level condition label for each cell.

After training the prediction model, River applies multiple attribution methods to determine the genes contributing to the model’s prediction behavior for the corresponding label in each input cell. After cell-level normalization, River provides multiple cell-wise gene scores to measure the relevance of each cell to its corresponding label in different aspects. River then combines the multiple cell-wise gene score to obtain a global summary statistic for each gene in the input cell population, resulting in a final rank for the input gene list for each attribution method. Finally, River adopts rank aggregation methods to combine the different ranks obtained by various attribution methods to produce a final gene rank.

### Alignment for multiple input slices

River requires spatial location coordinates as the predictive model input. When handling input slices from different spatial coordinate systems where the spatial locations have not been previously registered, the Spatial-Linked Alignment Tool (SLAT)^69^ is selected for its flexibility and scalability. In experiments involving two slices, a single SLAT alignment is performed.

For experiments with more than two slices, one slice is designated as the base slice, and SLAT alignment is then conducted for each of the remaining slices relative to this base slice. The outcome of the SLAT algorithm is a matching list, which identifies the corresponding cells of the remaining slices concerning the base slice. This matching list enables the projection of the remaining slices’ coordinates into the same spatial coordinate system as the base slice. Formally, given the input slice *K* with *n*_*k*_ cell/spots, and the corresponding spatial coordinate matrix *C*_*k*_, we choose slice 0 as the base slice. Thus, we will have the matching list obtained from SLAT for the remaining slice *m*_*k*_, and we have the new aligned coordinates for each input slice *k* as the input for River:

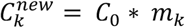

### Prediction model architecture

Given the multi-slice annotated dataset, the input of the River prediction model is composed of gene expression *x*_*i*_ for each cell and the corresponding label *y*_*i*_ as a one-hot vector for the cell of its belonging slice label, the aligned coordinates 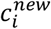. The prediction model encoder is composed of two parts: the gene expression encoder, which extracts the feature from *x*_*i*_, and the position encoder, which extracts the spatial information from 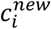. We utilize two MLPs as the position encoder *f*_*pos*_ and expression encoder *f*_*exp*_ separately. River adopts the double-branch architecture to encode the position information and the expression information separately and then combines them in the latent space to obtain the spatial-aware gene expression latent. We have the position latent vector 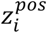:

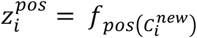

And the gene expression latent vector 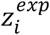:

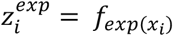

River then concatenates the two latent vectors to get the spatial-aware gene expression latent vector *Z*_*I*_ for the input cell:

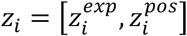

This latent vector is then sent into a following MLP classifier to get prediction logits 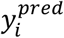:

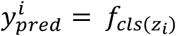

And the model is trained using the cross-entropy objective *L*_*ce*_ with the provided cell-level label *y*_*i*_:

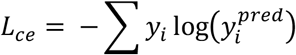

### Attribution methods

As aforementioned in the River framework composition, apart from the prediction model part, another important component of River is the attribution module. River’s ability to identify DSEP genes is based on the assumption that only genes with significant spatial expression pattern shifts across multiple slices can contribute to the prediction model’s ability to classify different slices. Thus, the attribution module aims to select the genes that contribute the most to the model’s decision process. In other words, River rank each gene using a post-hoc attribution method based on each gene’s spatial expression pattern.

River employs three state-of-the-art attribution methods: Integrated Gradients^70^, DeepLift^71^, and GradientShap^72^. River applies these three attribution methods to attribute the model’s prediction logits on ground-truth class *s*_*i*_(*x*_*i*_) back to the input features *x*_*i*_, yielding a weight vector 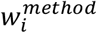 for each gene in each input cell that signifies the importance of each input gene.

One of the simplest attribution methods for deep learning is the Gradient * Input technique, initially proposed to enhance the clarity of attribution maps^71^. This method calculates attribution by partial derivatives of the output corresponds to the input and then multiplying these derivatives by the input. However, Gradient * Input is insufficient for handling complex scenarios. Therefore, we utilize three tailored attribution methods from modern deep learning to address these complexities.

For Integrated Gradient, we utilize 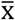 represents the baseline input. we choice the all zero gene expression input vector as the baseline in all the following methods, in cell *i* we have

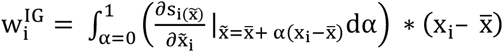

Integrated Gradient similarly to Gradient * Input, computes the partial derivatives of the output with respect to each input feature. However, while Gradient * Input computes a single derivative, Integrated Gradients computes the average gradient while the input varies along a linear path from a baseline 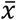 to *x*_*i*_.

As for GradientShap, it approximates SHAP values by computing the expectations of gradients by randomly sampling gradients since the exact SHAP value calculation is too expensive and cannot scale to large-scale cell datasets. In River, we apply the GradientShap method by adding white noise to each input gene expression n (n=5 in River default parameters) times,

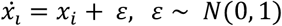

And then construct a random point *q*_*i*_ along the path between the baseline and the noisy input with scale parameter λ:

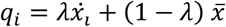

Then we compute the gradient of outputs with respect to those selected random points and get the final attribution score 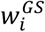:

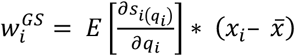

DeepLIFT is an attribution recursive prediction explanation method for neural networks that proceeds in a backward fashion. The importance for DeepLIFT is based on propagating activation differences on each neural unit in the neural network. Thus, compared with the previous two methods, DeepLIFT is proposed only for neural network attribution. For the sake of convenience, we utilize the modified chain rule notation introduced in Given two neural units (*a, b*) in the multi-layer perceptron, there must exist a path set *P* from *a* to *b* in the neural network due to the fully-connected property of the MLP. We can define a modified chain rule based on this path set *P*_*ab*_:

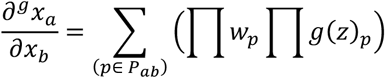

Where *z* indicates the linear transformation for each neural unit. For the neural input unit *j* and the output unit *i*, we have the output *z*_*j*_:

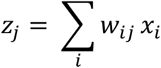

And *g* can be any other nonlinear transformation function. When *g* is the original non-linear activation function in the model, this modified chain rule will be equal to the partial differential.

With this notation, given the baseline and input gene expression, we have 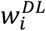:

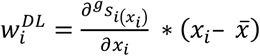

Where

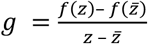

And *f* is the prediction model’s original activation function, (ReLU^73^ in our default setting), and 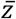 indicates the baseline corresponding *z*.

After multiple attributions, we can have the cell-level attribution score vector for each method. River normalizes them per cell-wise. The motivation here is to follow per-cell gene expression normalization preprocess, which can normalize different cells’ score on the same scale while maintaining heterogeneity. River normalizes each cell’s absolute weight vector using L2 normalization:

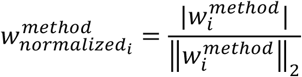

Then, River computes the global attribution vector for each method by averaging the normalized vectors:

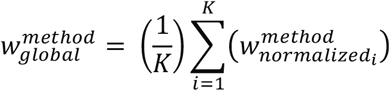

Finally, River ranks the genes based on their global attribution scores to determine their relative importance:

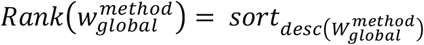

River further tries to aggregate these three different methods ranks into one final rank.

### Rank aggregation for multiple-attribution

Given the global weight vector for each gene derived from three state-of-the-art attribution methods (Integrated Gradient, DeepLIFT, and GradientShap), River aims to aggregate the attribution results for each method to obtain robust and stable attribution results since the three attributions indicate three different attribution perspectives (average gradient, estimated SHAP value, and neural unit activation). Here, each method provides a rank of genes based on their contribution to the prediction model’s predictions. To aggregate these rankings into a final comprehensive ranking, we employ the Borda count method^74^.

The procedure for applying the Borda count method to our context is as follows: for each attribution method, assign a score to each gene based on its rank. If a gene is ranked first, it receives a score equal to the total number of genes *N*; the second-ranked gene receives *N*−1, and so on, with the lowest-ranked gene receiving a score of 1. We then aggregate the scores for each gene across all attribution methods:

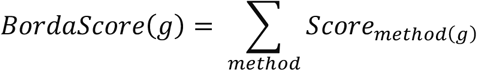

where *Score*_*method(g)*_ is the score assigned to gene *g* by the ranking from a particular method. Finally, the final rank of the genes can be obtained based on aggregated Borda scores:

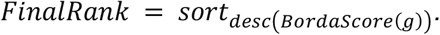

This Borda count aggregation method ensures that the final ranking reflects a balanced consensus across the different attribution methods, taking into account the unique perspectives each method offers on gene importance.

### Simulation dataset

#### Data generation

To generate the simulated dataset, we first created a control slice consisting of a square region with four distinct spatial domains using SRTsim^75^: domain A, domain B, domain C, and domain D. The gene expression in this slice was simulated using the SRTsim reference-free simulation procedure. Specifically, the simulated slice was composed of 980 randomly placed cells in a square shape. We initially divided the slice into domains A, B, C, and D with square shapes. SRTsim generated gene expression counts for 1100 genes by sampling from a ZINB distribution with the following parameters: zero percentage 0.05, dispersion 0.5, and mean value 2, and then randomly assigned them to 980 generated spatial locations in the square slice.

Furthermore, to mimic the true distribution on a real slice, SRTsim was applied to generate domain-specific differential genes for each given domain. After this procedure, the 1100 genes were split into three groups: higher signal genes (100 genes), lower signal genes (100 genes), and background genes (900 genes). Higher signal genes showed higher fold-changes in each domain with the domain-specific fold-change ratios (1.0 for domain A, 2.0 for domain B, 1.5 for domain C, and 3.0 for domain D). Meanwhile, the lower signal genes exhibited lower fold-changes in each domain with the same predefined domain-specific fold-change ratios. Finally, background genes maintained the same expression pattern as the original generation distribution.

We regarded this simulated slice as the control slice for our subsequent condition-perturbed slice generation. Here, we aimed to perturb the background genes to generate differential spatial expression patterns across the condition and control slices. The reason for not utilizing the signal genes was that the Spatially Variable Gene (SVG) property on the original slice would influence the perturbation efficiency. The signal genes acted as distractors to improve the benchmarking difficulty, as we did not modify the signal genes across the condition and control slices.

For the perturbation process, we first randomly chose 200 target genes from the background genes for each perturbation. The perturbation process consisted of two main parts. The first part involved the random permutation of the spatial locations for the chosen target gene expression values for each cell. For each pair of spatial location and target gene expression in each cell, we randomly permuted the corresponding relation. Each cell was assigned a new target gene expression value that originally belonged to another cell in the slice. This procedure ensured that only the spatial pattern of the chosen target genes was influenced, while the expression levels of other genes remained unchanged. In the second step, we altered the fold-change ratio for the chosen target genes only in specifically chosen domains by applying a twofold change on the permuted slice. We obtained the final version of the condition slice after this step. The two perturbation steps ensured that the spatial gene expression pattern of the chosen target genes in the condition slice differed from the control slice in both spatial distribution and gene expression value.

We generated our benchmarking dataset using two target gene sets and three chosen domains (domain B, domain C, domain D), resulting in a total of six simulated benchmarking datasets.

#### Implementation details of River

We introduce the implementation details of River for benchmark experiments and subsequent real data experiments.

For the benchmark dataset, because the condition slice is simulated based on the control slice, their spatial coordinates are located in the same space. Thus, pre-alignment of the two input slices is not needed. The prediction model part of River comprises a gene expression encoder, a position encoder, and the final classifier, forming three main parts. All of them are two-layer MLPs with the ReLU function as the activation function. The two encoders have the same hidden dimension of 64, while the classifier has a hidden dimension of 32. Dropout regularization is added during model training to avoid overfitting, with a dropout ratio of 0.3. For model training, the Adam optimizer is utilized with a commonly-used learning rate of 0.001 and a weight decay rate of 0.0001. The model is trained for 100 epochs, and the last epoch model is used as the attributed model. The batch size is set to 4096 to ensure efficiency and fast convergence. For the attribution part, the captum package is adopted to implement the three attribution methods, utilizing the default parameters of the official package to obtain stable attribution results. River selects the top-200 ranked genes, corresponding to the predefined target gene number in the benchmark experiment.

For other real data experiments requiring pre-alignment, SLAT is utilized as the pre-alignment tool. For spatial transcriptomics data, the PCA value of the gene expression is used as the input for SLAT, and the raw profiled expression value of the spatial proteomics is used as input due to its relatively lower dimension (less than 100). After choosing a base slice for each experiment, every slice is subsampled to the same cell number as the base slice to ensure each cell has a corresponding aligned coordinate in the base slice. The neighborhood graph is then constructed using the K-nearest cells (K=20), and the preprocessed data is sent into SLAT to get the aligned coordinates for non-base slices. Apart from the alignment procedure, real data experiments use the same model parameters as mentioned in the benchmark dataset section.

#### Competing methods

In this study, we compared three main categories of methods to identify the differential spatial expression patterns between condition and control slices in six different simulation benchmark datasets.

The first category of methods includes the previously highly variable genes (HVGs) selection methods. We adopted the most commonly used HVGs selection methods - Seurat, Seurat v3, and CellRanger as the baseline methods. In each experiment, given the condition and control slice gene expression as input, we defined the selected gene number as 200 (equal to the target gene number). The implementation of these three methods utilized the scanpy^24^ package with the default parameters.

The second category includes the conventional Spatially Variable Genes (SVGs) selection methods. However, as mentioned previously, these SVGs methods cannot handle multi-slice input. Therefore, modifications were made to these methods to enable them to perform the same task. For the significance-based methods, due to the difficulty in comparing the output significant statistics among slices, we utilized the absolute difference value (to ensure non-negative input) of the same position cells in two slices as the input for the following test-based SVGs methods. The motivation here is that the spatial variance of the difference value can reflect the spatial expression pattern to an extent. We utilized four commonly used methods: SPARKX, SpatialDE, Moran’s I, and Geary’s C as the test-based SVGs method baseline. Default parameters were utilized for both methods, and significant genes (p-value < 0.05) were regarded as detected positive genes, with the remaining as negative.

Apart from the test-based SVGs methods, there is another type of SVGs method: score-based SVGs detection. This method provides a score for each gene as output, indicating the spatially related situation for each gene, making comparison between slices possible. Here, we utilized Sepal as the baseline method in the score-based SVGs methods. The original code in the Sepal package was used. Transformation was applied to convert our input slice into spot-level data like Visium, since Sepal only accepts such format input. The official transformation function provided in Sepal was used to convert the slice into a spot-like arrangement, and Sepal was then applied to the two control and condition slices independently. The output scores were normalized into the range [0,1], making the scores among the two slices comparable. The absolute difference value for each gene was then calculated, and the spatial pattern change was ranked by this absolute difference score, with a higher value indicating a larger difference. As with the previous HVGs methods, a selected gene number k was set to select the top-k highest score genes, using k=200 as with HVG methods.

The final category comparison method is the 3D SVGs identification methods. These methods aim to find the genes which show significant spatial expression patterns considering 3-dimensional spatial information. We utilized the recently proposed BSP as our 3D SVGs baseline. The control and condition slices were stacked to obtain a pseudo-3D slice dataset, setting the z-coordinate for all control slices as 0 and condition slices as 1. The official version of BSP was then applied to this pseudo-3D input slice with its predefined 3D format. The output of BSP is also the significant statistics. Thus, significant genes (p-value < 0.05) were regarded as positive genes, with the remaining as negative genes.

### Evaluation metrics

We utilized the F1-score as the metric to measure each model’s performance in identifying Differential Spatial Expression Pattern (DSEP) genes. The reason for not adopting recall and precision as additional metrics is that, for the k selected methods (e.g., River, Sepal, and HVGs methods), the F1-scores are the same as recall and precision. Regarding the perturbed target genes as the positively labeled genes and the method selecting positive genes, we have the F1-score:

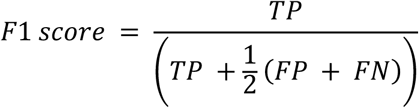

Furthermore, to evaluate the robustness of the k selected methods, the performance among k selected methods is compared by the F1 score on different k values.

### Analysis of the Stereo-seq mouse embryo dataset on batch-related genes

#### Dataset overview

We utilized the Stereo-seq mouse embryo multi-slice dataset, which is composed of eight different development stages at near single-cell resolution. Each development stage consists of different depth 3D slices from the same replicates. We utilized the E15.5 development stage as the input multi-slice dataset for River. This developmental time point slice is composed of four continuous depth slices along the z-axis, similar to the E16.5 development stage. We used the depth for each slice as the slice-level label and selected the slice with the lowest depth as the base slice, with 10,000 subsampled cells for each slice for alignment. River then applied fitting and selection on the input depth-informed cells dataset.

#### Slice integration and evaluation

In the visualization of the River-selected top-ranked genes, we observed that the top-ranked genes are highly related to the depth for input cells. In other words, it is possible to integrate the input cells from different batches to the same depth by using such depth information-preserved genes as input features. We used only the River selected top-20 genes as input to conduct the integration across two development stages (E15.5 and E16.5). It is worth noting that River did not see any E16.5 cells during training. Therefore, this integration not only supports that depth information-preserved genes can help integrate cells at the same depth but also provides evidence of the River-selected genes’ generalization. We utilized Harmony as our integration method due to its efficiency and accuracy and then evaluated the integration results using the scib package. This package first conducts a fine-grained search for the best clustering resolution of Leiden clustering and then utilizes the best cluster outcome to evaluate both biological-preserved metrics (Normalized Mutual Information (NMI), Adjusted Rand Index (ARI)) and batch-removal metrics (Graph Connectivity (GC), Integration LISI (iLISI) graph score). Specifically, we utilized depth as the biological information variable and the different development stages (E15.5, E16.5) as the batch variable.

NMI compares the overlap of two clustering:

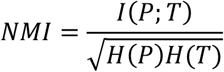

Where *P* and *T* are categorical distributions for the predicted and real clustering, *I* is the mutual entropy, and *H* is the Shannon entropy.

As for ARI, it considers both correct clustering overlaps while also counting correct disagreements between two clustering:

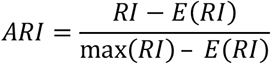

Where Rand Index (RI) computes a similarity score between two clustering assignments by considering matched and unmatched assignment pairs. Both of them evaluate whether the integrated embedding can capture biological information properly.

As for the batch-removal metrics, iLISI and cLISI are adopted from the Local Inverse Simpson’s Index (LISI) for the batch-related and biological preservation metrics. Here, we utilize the iLISI modified version in scib.

LISI scores are computed from neighborhood lists per node from integrated kNN graphs. Specifically, the inverse Simpson’s index is used to determine the number of cells that can be drawn from a neighbor list before one batch is observed twice. Thus, LISI scores range from 1 to *B*, where *B* is the total number of batches in the dataset, indicating perfect separation and perfect mixing, respectively, and scib rescales them to the range of 0 to 1. Given the total batch label-based LISI score set *X*, we have:

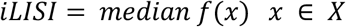

Where *x* indicates the previous LISI score for each batch label, we have

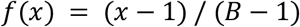

With higher iLISI indicating a better batch mix situation.

For cLISI, we need to modify the *f*(*x*) into *g*(*x*):

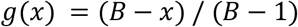

Where *B* here indicates the total cell type number and *x* indicates the LISI score for each cell type, and *X* is the total cell type-based LISI score set. We have:

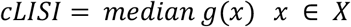

The GC metric quantifies the connectivity of the subgraph per cell type label. The final score is the average for all cell type labels *C* according to the equation:

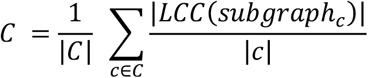

Where |*LCC*(*subgraph*_*c*_)| stands for all cells in the largest connected component in the dataset, and |*c*| stands for all cell numbers of cell type *c*.

These metrics examine the integrated embedding’s batch information mixture situation. Higher values indicate better batch removal performance.

### Analysis of the Stereo-seq mouse embryo dataset on development-related genes

#### Dataset overview

We utilized the Stereo-seq mouse embryo multi-slice dataset, which is composed of eight different development stages at near single-cell resolution. Each development stage is composed of different depth 3D slices on the same replicates. We utilized the eight development stages at the same depth as the input multi-slice dataset for River. The input multi-slice dataset is composed of [E9.5, E10.5, E11.5, E12.5, E13.5, E14.5, E15.5, E16.5] eight slices in the first layer for each time point 3D slice. We used the time point for each slice as the slice level label and selected the slice with the E9.5 as the base slice with 5000 subsampled cells for each slice for alignment. River then applied fitting and selection on the input development-informed cells dataset.

#### Pairwise silhouette score

To measure whether distance biological information preserves the condition in different input gene panel situations, i.e., whether the cell’s distance in the development timeline can be reflected in the latent space distance acquired from different input genes, we calculated the silhouette score pairwise for each cell pair of development stages.

Given any two development stage *p* and *q* pair, we have pairwise silhouette score *S*(*p, q*):

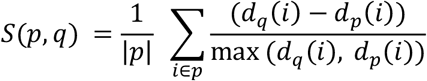

Where |*p*| indicates the cell number in p development stage and *d*_*p*_(*i*) denotes the mean L1 distance of cell i to all cell in distance p of input gene expression:

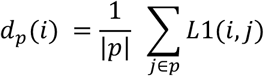

A smaller silhouette score indicates a shorter distance between two groups of cells in the gene expression space. After calculating the silhouette score for each development pair combination, hierarchical clustering can be conducted on the silhouette score vector for each development stage to determine the similarity situation for the development stage in the gene expression space.

#### Embedding evaluation based clustering

To evaluate the latent space’s embedding quantitatively, we adopted the scib^37^ pipeline to conduct the Leiden clustering with the best resolution search for each input gene set. The resolution search is in the (0.0, 1.0) range with 0.1 increasing each resolution. Then the embedding for each gene set is compared by the best NMI, ARI, and Cell-type LISI (cLISI) score values in the previous search pipeline. Higher NMI and ARI indicate better development stage information preservation and less noise information content for the input gene set

#### Binarize spatial gene expression

To evaluate River’s capability to identify pure spatial pattern shift genes across slices, we binarized the input gene expression data when fitting the prediction model (after the pre-alignment process). Specifically, for all input gene expressions, values greater than 0 were transformed into 1, while 0 values remained unchanged. This binarization process removes the influence of gene expression values and preserves only the spatial expression patterns, which we refer to as the pure spatial pattern. Apart from this binarization process, other parameters and settings remained the same as in the normal format.

After identifying the top-k binarized pure spatial pattern shift genes, we compared them with the previous top-k expression value-informed selected genes. We then selected the genes found only in the pure spatial pattern shift list as the unique pure spatial pattern shift genes for downstream analysis.

#### Gene set enrichment analysis

We conducted gene set enrichment analysis for the top-50 unique pure spatial pattern shift genes using the Enrichr API in the gseapy Python package, with GO Biological Process 2023 as the reference gene set. The cutoff for significantly enriched gene sets was an FDR-adjusted p-value of less than 0.05.

### Analysis of diabetes-induced and WT mouse testis

#### Dataset Overview

We applied River to disease-related spatial transcriptomics multi-slice datasets to demonstrate the application diversity of River. The dataset is composed of six Slide-seq slices from three leptin-deficient diabetic mice (Diabetes) and three matching wild-type (WT) mice. The WT-1 sample was selected as the base slice, and each slice was subsampled into 10,000 cells for alignment. The Diabetes and WT phenotypes for each slice were used as the corresponding slice labels.

#### Multi-slice spatial co-clustering

We applied CellCharter to the top-200 genes selected by River to achieve consistent spatial domain clustering on the input slices. CellCharter is a deep learning-based method that incorporates single-cell dimension reduction techniques to remove batch effects among slices and utilizes spatial coordinates to form a spatial graph. It then conducts automatic resolution-selected clustering based on a batch-integrated embedding spatial graph to obtain consistent spatial domain clustering across multiple input slices.

Following the CellCharter official tutorial for spatial transcriptomics data, we used the top-200 genes selected by River as input, employing scVI^76^ for dimension reduction with default parameters. We chose a clustering resolution range from 2 to 10, performing the clustering process 10 times with CellCharter’s AutoK process to ensure robust results.

#### Gene set enrichment analysis

We conducted gene set enrichment analysis for the top-50 DSEP genes using the Enrichr API in the gseapy Python package, with KEGG, Jensen TISSUES, and Elsevier Pathway Collection as reference gene sets. The cutoff for significantly enriched gene sets was an FDR-adjusted p-value of less than 0.05.

### Analysis of the human TNBC MIBI and mouse lupus CODEX dataset

#### Dataset Overview

We applied River on two disease-related spatial proteomics multi-slice datasets to show the modality-agnostic property of River and its further potential in clinical applications. The first disease spatial proteomics dataset is the human Triple Negative Breast Cancer (TNBC) MIBI dataset. The dataset is composed of 41 slices with three different TNBC subtypes featured by the immune cell infiltration condition: 15 Mixed (high immune infiltration) slices, 19 Compartmentalized (distinct tumor and immune cell regions) slices, and 5 Cold (low immune cell presence) slices. We randomly chose one slice from each subtype as the fitting multi-slice dataset for River and the remaining 38 slices as the hold-out slices. The chosen Mixed slice was regarded as the base slice, and every slice was subsampled into 2000 cells to conduct alignment. The second spatial proteomics multi-slice dataset is composed of nine CODEX mouse lupus spleen samples (3 WT and 6 lupus mouse samples). We selected one slice in each condition randomly and preserved the remaining 6 slices as the hold-out slices. The WT slice was regarded as the base slice and subsampled into 20,000 cells in each slice for alignment.

#### Panel Generalization Evaluation

In the disease-related proteomics multi-slice dataset, to evaluate the River-selected panel generalization and clinic application potential, it is assumed that the River selected top panel can be utilized in the unseen slice phenotype identification. The mean value of the selected panel expression on the slice is utilized as the input feature, and the predictive power of the panel feature is evaluated by conducting 5-fold cross-validation on hold-out slices in the original dataset with different baseline classifiers. We utilized the default parameters for three commonly used models (Support Vector Classifier (SVC), Logistic Regression (LR), and Random Forest (RF)) as the baseline classifiers. The 5-fold accuracy is compared for different panel selected situations (different top-k [2, 4, 6, 8, 10]) of River selected panel, randomly selected panel, and full panel.

#### Computational resources

All experiments were performed on a server running Ubuntu 22.04 with a 32-core Intel(R) Xeon(R) Gold 6338 CPU @ 2.00GHz and an Nvidia A800 (80G) GPU.

## Data availability

The Stereo-seq mouse embryo development dataset^33^ can be obtained from: https://db.cngb.org/search/project/CNP0001543

The Slide-seq mouse diabetes dataset^48^ can be obtained from: https://www.dropbox.com/s/ygzpj0d0oh67br0/Testis_Slideseq_Data.zip?dl=0

The CODEX mouse lupus dataset^26^ can be obtained from: https://data.mendeley.com/datasets/zjnpwh8m5b/1

The MIBI human TNBC dataset^52^ can be obtained from: https://mibi-share.ionpath.com

The MERSCOPE mouse brain dataset can be obtained from: https://vizgen.com/resources/using-merscope-to-generate-a-cell-atlas-of-the-mouse-brain-that-includes-lowly-expressed-genes/

## Code availability

The Python implementation and tutorial of River is available at https://github.com/C0nc/River.

## Acknowledgments

Z.Y. acknowledges the support by National Key R*i*D Program of China (2023YFF1204800), National Nature Science Foundation of China (62303119), Shanghai Science and Technology Development Funds (23YF1403000), Chenguang Program of Shanghai Education Development Foundation and Shanghai Municipal Education Commission (22CGA02), and Shanghai Science and Technology Commission Program (23JS1410100).

## Author contributions

Z.Y. and Y.C. conceived and designed the study, developed the computational methods, performed the analysis, and wrote the manuscript.

## Inclusion *i* Ethics

Not relevant.

---

**Supplementary Fig. 1.**
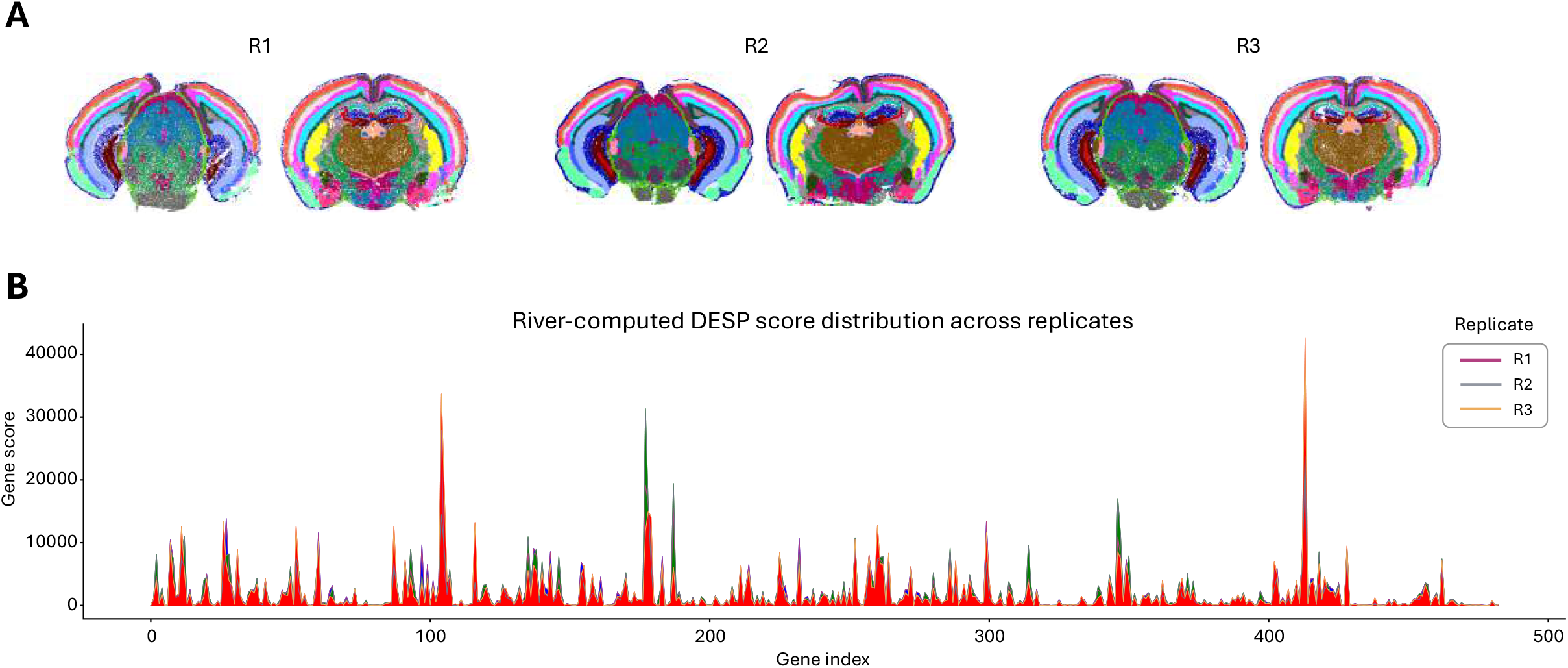
Scalability and reproducibility of River on MERSCOPE mouse brain dataset. **A**, Dataset: MERSCOPE mouse brain dataset composed of three replicates with two sectioning positions in the whole mouse brain, each slice containing > 70,000 cells. **B**, The River score (IG) distribution for each gene in three replicates experiment, showing high consistency.

**Supplementary Fig. 2.**
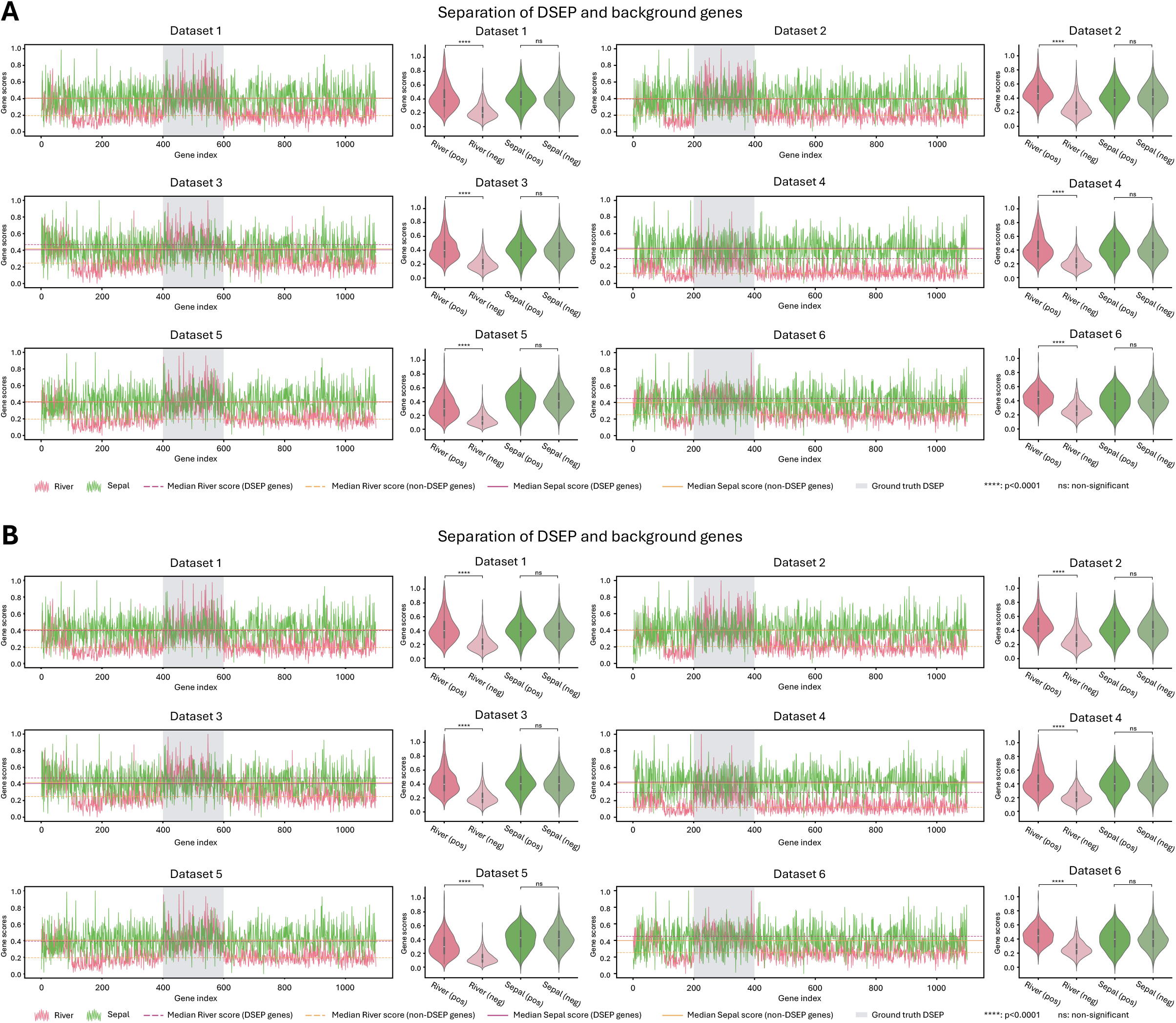
Separation of DSEP and background genes using River score (associated with Fig. 2E). **A**, Comparison of score distribution between River and Sepal. River’s attribution method is DeepLift. For each dataset, the left line chart indicates the score value for each gene, where positive genes (Ground truth DSEP genes) are expected to obtain larger scores compared with remaining negative genes. The right violin plot indicates the score distribution for the two methods between the DSEP and non-DESP genes. P-values are obtained using rank-sum test. **B**, Comparison of score distribution between River and Sepal. River’s attribution method is GradientShap. For each dataset, the left line chart indicates the score value for each gene, where positive genes (Ground truth DSEP genes) are expected to obtain larger scores compared with remaining negative genes. The right violin plot indicates the score distribution for the two methods between the DSEP and non-DESP genes. P-values are obtained using rank-sum test.

**Supplementary Fig. 3.**
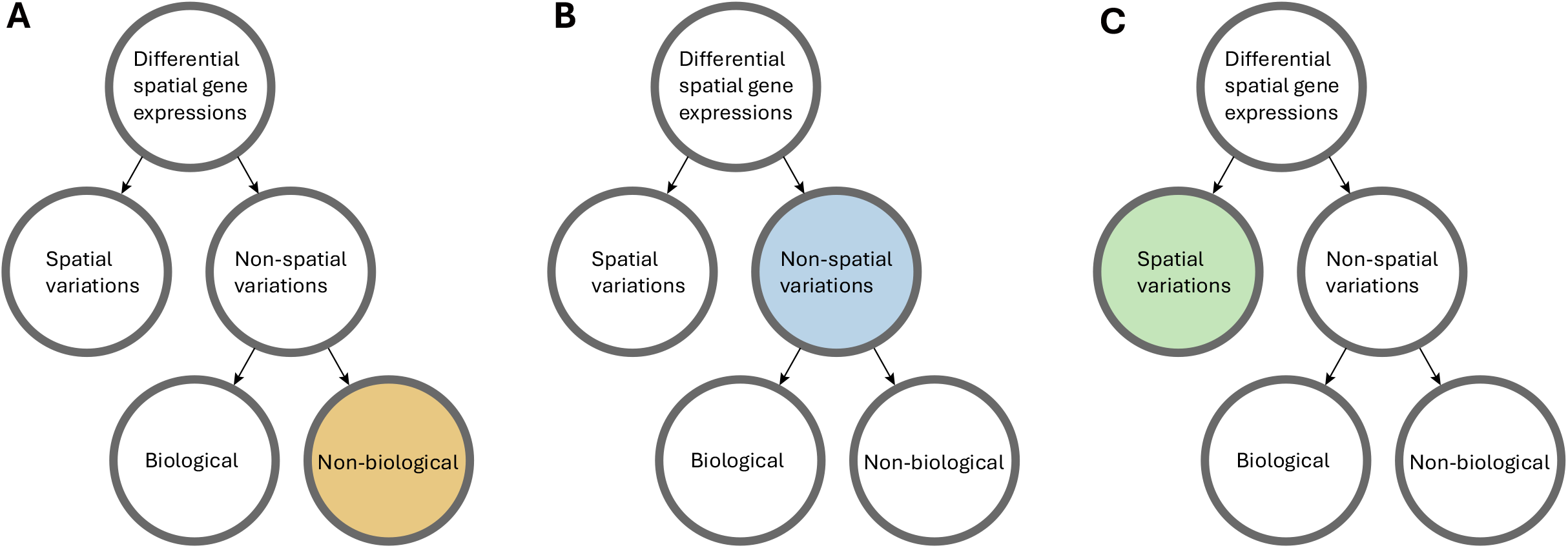
Factors contributing to differential spatial gene expressions. **A**, Associated with Fig. 3, where non-biology signals is the major variation. **B**, Associated with Fig. 4B-F, where the non-spatial signals is the major variation. **C**, Associated with Fig. 4G-H, where the spatial signals is the major variation.

**Supplementary Fig. 4.**
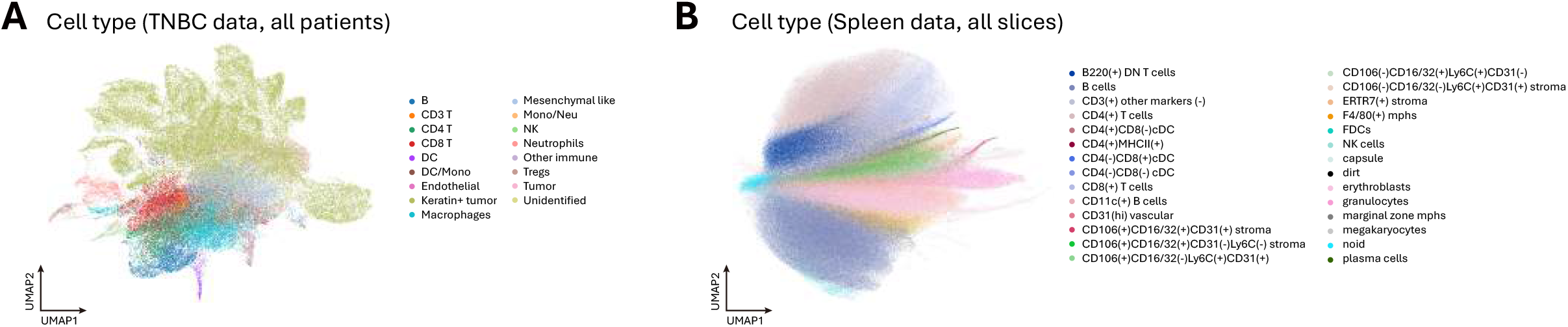
UMAP of all cells in spatial proteomics datasets. **A**, The TNBC MIBI-TOF dataset. **B**, The spleen CODEX dataset. Cells are labeled according to cell types in original papers.

